# New whole slide image analysis method for detection and quantification of hepatocellular vacuolization in European flounder (*Platichthys flesus*) in multipollutant exposure context

**DOI:** 10.64898/2026.10.08.757637

**Authors:** Valentin Geslin, Mamadou Diop, Jérôme Couteau, Rachid Amara, Thor Ole Gulsrud, Jean Laroche

## Abstract

A new whole slide image analysis method was developed for hepatocellular vacuolization detection and quantification in European flounder (*Platichthys flesus*) liver. Histopathological quantification allows a better understanding of how contaminants and biotic factors might influence lesion pathogenesis in a multipollutant exposure context. One of the main findings is the significant difference in the extent of the pathology between the polluted and the control sites, with *P. flesus* sampled in the Seine estuary presenting a higher proportion of tissue affected by hepatocellular vacuolation (6.88 ± 3.01%) compared to fish sampled in the Canche estuary (5.46 ± 2.86%). This result is coherent with the previous qualitative histological assessment done on the same histological samples, it helps validate the method and brings new insight into the pathology intensity level with the level of contamination on each site. The other main finding is the potential synergetic effect of the pollutant mixture on the pathology. The multipollutant analysis showed a positive mixture effect on the proportion of tissue affected by hepatocellular vacuolation in fish from the Seine estuary, while this effect is not observed in fish sampled in the Canche estuary. This study highlights the benefits of developing novel bio-image analysis methods for environmental research. Histopathological diagnostic quantification allows a better understanding of the potential factors inducing and exacerbating the pathology, as well as the potential mixture effect in a context of multipollutant contamination.

## Introduction

Hepatocyte vacuolization in flatfish species is a well-known pathological condition where excessive lipid accumulates in cytoplasmic vacuoles of hepatocytes. Pathological evidence of the disease can be difficult to differentiate from normal accumulation of lipid however, affected cells usually contain larger lipid droplets or numerous small vesicles (Vethaak et al., 1996). In flatfish species the disease is triggered by hepatic metabolism disruption that has been linked to both endogenic and exogenic factors, among them, exposure to pollutants is one of the most documented (Bernet et al., 1999; Feist et al., 2004; Lang et al., 2017; Moore et al., 1997; Stentiford et al., 2003; Vethaak et al., 1996; Vethaak et al., 2009). Excessive cytoplasmic lipid accumulation is generally regarded as an early non-neoplastic toxicopathic lesion, and its relevance as a histopathological biomarker for xenobiotic compound exposure in flatfish species is well established (Feist et al., 2004). Hepatocyte vacuolization diagnosis in *Platichthys flesus* is commonly achieved through histopathological examination of liver tissue. The lesion is qualitatively graded, based on its severity and spread within the tissue (Feist et al., 2004; Stentiford et al., 2003). Although qualitative assessment can be informative, it remains poorly characterized without quantitative measurements (Wolf and Wheeler, 2018). As previously mentioned, hepatocyte vacuolization in *P. flesus* has been linked to exposure to contaminants, but its pathogenesis remains poorly understood in a multipollutant exposure context (Köhler, 1990; Köhler et al., 2002; Stehr et al., 1998). A multipollutant exposure analysis addresses the independent effects of each contaminant, the overall effects of multipollutant mixtures, and potential synergetic or antagonistic effects of the mixture on a specific biomarker (Zhu et al., 2024). Since recent advances in bio-image analysis allows to detect and quantify histological pathologies more accurately, multi-pollutant exposure analysis can now be used with histopathological biomarkers (Jain et al., 2024; Kong et al., 2011; Mulrane et al., 2008). This article presents a new whole slide image (WSI) analysis method to detect and quantify the spread of hepatocyte vacuolization in *P. flesus* liver. A multipollutant exposure analysis was used to characterize potential relationships between this disease and the multiple pollutants measured in the same fish.

## Material and methods

### Original data set

Data originated from the following article: *Impacts of chemical stress, season, and climate change on the flounder population of the highly anthropised Seine estuary* (Laurent et al., 2022) (https://doi.org/10.1007/s1022-20000-y). One of the goal of the original project was to assess the impact of Seine estuary water quality on *P. flesus* fitness. A multi-biomarker approach coupled with multipollutant measurement in fish liver was used for a comprehensive assessment of pollution effect (Laurent et al., 2022). Canche estuary was used as a reference site because the river and its estuary are relatively preserved from polluting industry compared to the Seine river. A total of respectively 88 and 53 adult fish were sampled in the Seine and Canche estuaries between 2018 and 2019. The first sampling was conducted during the reproductive period (winter 2018), and a second sampling occurred during the inter-reproductive period (summer 2019). Similar effort were made to sample female and male *P. flesus* evenly. However, the number of fish sampled in the two stations, at the different seasons and between sexes are not equal, each group have a different sample size (Table 1). Moreover, some biomarker analysis and pollutant measurement were missing from the original dataset. For this article, only fitness index, reproductive markers, liver lipid content, and contaminant data from the original article were used.

**Table 1.** Number of sample and mean total body length between station, season, and sex.

| Station | Season | Sex | Sample size | Mean body length [min-max](cm) |
| --- | --- | --- | --- | --- |
| Canche | Summer | F | 16 | 22.1 [16.9-29.2] |
| Canche | Summer | M | 6 | 23.4 [21.4-26] |
| Canche | Winter | F | 18 | 30.0 [27.2-33.7] |
| Canche | Winter | M | 13 | 26.6 [23.5-30.2] |
| Seine | Summer | F | 35 | 30.9 [24-38.5] |
| Seine | Summer | M | 23 | 30.1 [25-37.6] |
| Seine | Winter | F | 19 | 32.4 [26.7-44.1] |
| Seine | Winter | M | 11 | 30.4 [25.1-34.7] |

### Fitness index

Fulton’s condition factor (K) (1) is a measure of the relationship between fish length (L) and body weight (W).

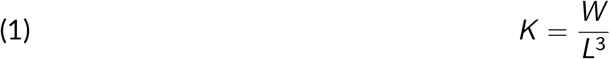

Hepatosomatic index (HSI) (2) measure the relationship between total fish weight (W) and liver weight (WL).

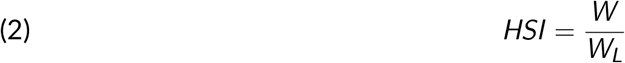

The lipid storage index is based on the ratio of the quantity of triglyceride (TG) and free sterol (FS) measured in muscle (3). These indexes are considered as indicators for fish’s health and nutritional status (Amara et al., 2007; Peig and Green, 2010; Wuenschel et al., 2019).

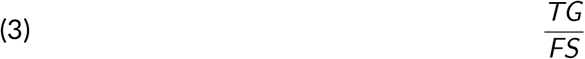

Growth rate was determined by back-calculating otolith growth rings (Laroche et al., 2017). Age was extrapolated from total body length according to Von Bertalanffy growth equation (4), where *α* is age, k is the growth coefficient, *t*_0_ is the theoretical age when size is zero, and *L*_*∞*_ is the theoretical age when size is zero.

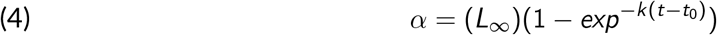

### Reproductive markers

Female reproductive markers were: plasmatic estradiol level (E2), gonadosomatic index (GSI) (5), which measures the relationship between gonad weight (*W*_*L*_) and body weight (W).

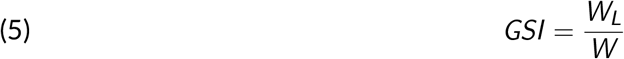

Sexual maturation index (SMI) (6) reflects individual maturation stage. It is calculated with number of oocytes at stage 1 (*N*_1_), number of oocytes at stage 2 (*N*_2_), number of oocytes at stage 3 (*N*_3_), number of oocytes at stage 4 (*N*_4_) and total number of analyzed oocytes (*N*_*t*_).

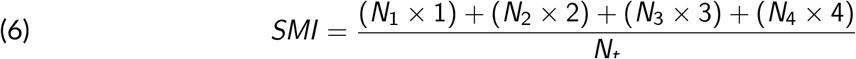

Males reproductive markers were: comet assay on spermatozoid, level of plasmatic steroid hormones like 11-ketotestostérone (11KT), vitellogenin (VTG) levels, and GSI (4).

### Liver lipids content

Six neutral lipid classes (sterol esters (SE), glyceride ethers (GE), triacylglycerols (TG), free fatty acids (FFA), fatty alcohols (ALC), and free sterols (FS)) and seven polar lipid classes (sphingomyelin (SPG), lysophosphatidylcholine (LPC), phosphatidylcholine (PC), phosphatidylserine (PS), phosphatidylinositol (PI), cardiolipin (CL) and phosphatidylethanolamine (PE)) were measured in *P. flesus* muscle. Neutral lipids (except FS) are storage lipids, whereas polar lipids and FST are associated with membrane lipids.

### Liver histopathology

Histopathological lesions were categorized according to Feist et al., 2004. Contrary to what was done in the original article, histopathological symptoms were not graded but analyzed through their prevalence in each group.

### Contaminants

Levels of different pollutants were measured in fish liver: polycyclic aromatic hydrocarbon (PAH) (naphthalene, dibenzothiophene, fluoranthene, pyrene, and chrysene), polychlorinated biphenyls (PCB), hydroxypyrene, pesticides (4’4 DDE, 4’4 DDD), polybrominated diphenyl ethers (PBDE), and different metal (Ag, Al, Cd, Co, Cr, Cu, Fe, Hg, Mg, Mn, Mo, Ni, Pb, Sr, Ti, V, and Zn) were measured in fish liver.

### Image analysis

Histological slides were scanned using a Leica GT450 scanner. QuPath (0.5.1), Python (3.13.3), and Fiji (2.16.0) were used for image analysis. WSI analysis method consists of six main steps, including tissue detection, tiling, tile sorting, color normalization, detection and quantification, and results compilation. The different steps are described in detail in the following sections.

#### Tissue detection

Tissue regions were identified using a thresholding method. This operation identifies pixels with values above the specified threshold and pixels with values below, allowing discrimination of tissue from the white background. A pixel value threshold of 220 was selected, with minimum tissue area of 1 mm^2^, and annotations touching image boundaries were excluded. Tissue areas were exported for downstream processing as well as pixel dimensions from each WSI.

#### Tiling

Tiles of 1600 µm^2^, with a down-sampling factor of 1, were generated from WSI regions where tissue was previously detected. Tiles were exported as raster graphics files (.png). For each tile, a twin tile representing only pixels where tissue was detected was exported in parallel.

#### Tile sorting

Tiles were sorted to remove tissue edges, histological structure (pancreatic tissue, liver tubule, mural plates, bile ducts, blood vessels…), and artifacts (tissue fracture, incision, puncture… ) as they could produce false positives, and affect the analysis outcome. Tile sorting consists of counting non-white pixels on each tile’s twin. Non-white pixels supposedly represent tissue area in each tile. If the ratio of non-white pixels was higher than 90% of total pixels in the image, it was selected for further analysis. In other words, only tiles with at least 90% of tissue were selected for further analysis.

#### Color normalization

Color normalization is used to homogenize colors before image analysis. Variation in image colors can originate from sample preparation or imaging conditions. Selected tiles were normalized using unmixing (separation of the H&E stains) and appearance normalization as described in Macenko et al., 2009.

#### Hepatocellular vacuolization detection and quantification

Each tile was processed using a HSB color threshold to isolate pale regions from the rest of the tissue. A first mask of this color detection was produced. Edge detection and particle analysis are then used on the mask. Particle analysis detects objects between 5-200µm^2^ with a circularity index between 0.30 and 1.00. The area of the detected objects were exported as results.

#### Results compilation

Results across all processed tiles of each image were aggregated, and the percentage of tissue area affected by hepatocellular vacuolization (PTAHV) was calculated for each image based on the total tissue area detected and the number of selected tiles used for the analysis.

### Relationships with other biomarkers and contaminants

The relationships between PTAHV and fitness index, reproductive marker, liver lipids content, histopathological lesions, and contaminant were then analyzed. Different models (linear regression, cubic polynomial regression, quadratic polynomial regression, and generalized additive model), both linear and non-linear, were compared to search for any significant relationships between PTAHV and every individual factor. The relationship between PTAHV and each single pollutant was assessed following the same method. Multipollutant exposure analysis was performed using a Bayesian Kernel Machine Regression (BKMR) model:

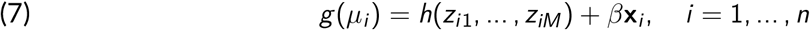

Where *g* is a monotonic link function, *µ*_*i*_ = *E* (*Y*_*i*_ ), *h* is a flexible function of the predictor variables *z*_*i*1_, …, *z*_*iM*_, *x* is a vector of covariates assumed to have a linear relationship with the outcome, and *β* is the corresponding vector of coefficients. The predictors *z* referred to exposure variables and *h*(*·*) as the exposure-response function represented with a Gaussian kernel machine regression expressed as:

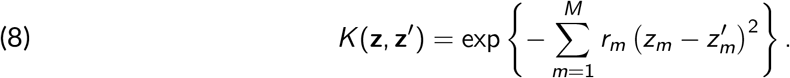

*z* and *z*^*′*^ represent vectors of predictors for two different individuals, and *r*_*m*_ *≥* 0 denotes the tuning parameter that controls the smoothness of *h* as a function of the exposure *z*_*m*_. PTAHV variations between individuals were analyzed, taking into consideration all contaminant levels, potential interactions between pollutants, and covariates (station, season, and sex).

### Statistical analysis

Statistical analysis was performed with R (4.5.2). A 0.05 significance level was used for all statistical tests. When homoscedasticity and homogeneity of variances were respected, a twoway ANOVA type III was used to test differences between groups and to account for unbalanced design. It was followed by a post-hoc Tuquey test. Otherwise, Kruskal–Wallis test was applied, followed by a post-hoc Dunn’s test. Multipollutant exposure analysis was performed with the Bayesian Kernel Machine Regression (BKMR) package (Bobb et al., 2015).

### Method validation

The validation protocol assesses the WSI analysis accuracy and its reliability in terms of false positives and false negatives. A subsample of 363 tiles was randomly drawn from all selected tiles from all WSI. The subsample represents about 1% of the total number of selected tiles (9413 selected tiles in total). Tissue area affected by hepatocellular vacuolization was manually annotated on each tile with QuPath. Areas obtained by manual annotations were compared to the area detected with the WSI analysis method, as well as the overlapping area between the two methods.

## Results

### Method validation

No significant differences (Wilcoxon rank-sum test, p > 0.05) were observed between the mean PTAHV detected by manual annotation and WSI analysis. Overall mean percentage of overlapping area was 72.5% (± 19.2%).

### Unbalanced sampling & Cofounding factor

A significant difference (Anova type III p < 0.05) in fish’s mean total body length between station, season, and sex was observed (Figure 1). *P. flesus* from Seine were significantly longer (31.0 ± 4.09 cm) compared to those from Canche (26.0 ± 4.15 cm). Fish from Canche present a significant difference (Anova type III, p < 0.05) in mean total body length between seasons, with fish sampled in summer being significantly smaller (22.5 ± 3.24 cm) than fish sampled during winter (28.6 ± 2.57 cm), while this difference was not observed in Seine. No significant difference (Anova type III, p > 0.05) in mean total body length between sexes. This result suggests that fish from the two stations may not have the exact same age; therefore, it was integrated as a cofounding factor into the statistical analysis.

**Figure 1.**
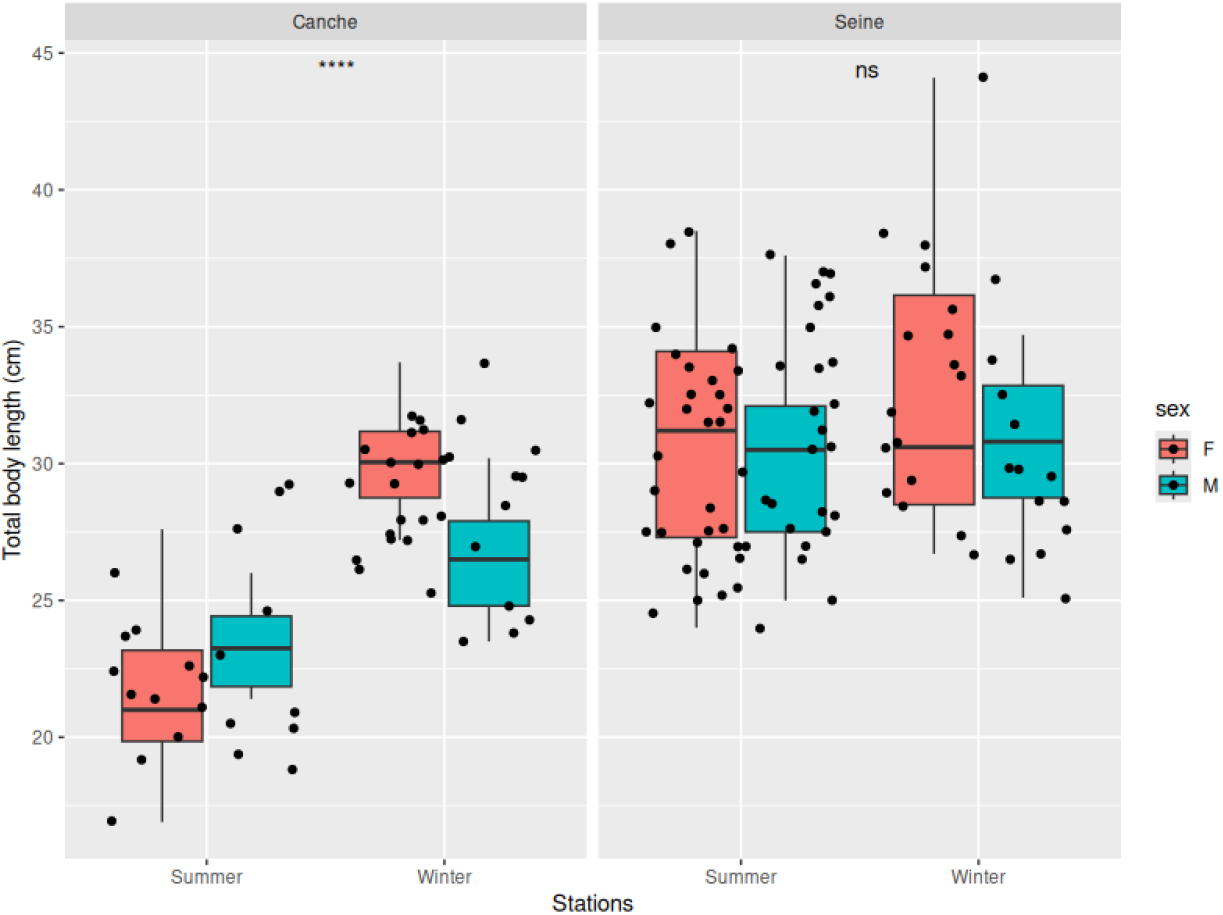
Box plot representing *P. flesus* total body length between station, season, and sex. Fish from Canche present a significant difference in mean total body length between season, with fish sampled in summer being significantly smaller (22.5 ± 3.24 cm) than fish sampled in winter (28.6 ± 2.57 cm), while no significant difference was observed between season for fish sampled in Seine. Fish from Seine were significantly longer (31.0 ± 4.09 cm) compared to those from Canche (26.0 ± 4.15 cm). No significant difference in mean total body length between sexes.

The proportion of selected tissue area per WSI was calculated by dividing selected tissue area (µm^2^) multiplied by 100, by total tissue area (µm^2^). It ranges from 89.2% to 29.1% with a mean of 72.9% (± 15%). A significant negative correlation (cor = -0.23, p < 0.05) was found between the proportion of selected tissue area and PTAHV (Figure 2). This results suggest that the proportion of selected tissue area might negatively affect the outcome of the WSI analysis, therefore it was integrated as a cofounding factor into the statistical analysis.

**Figure 2.**
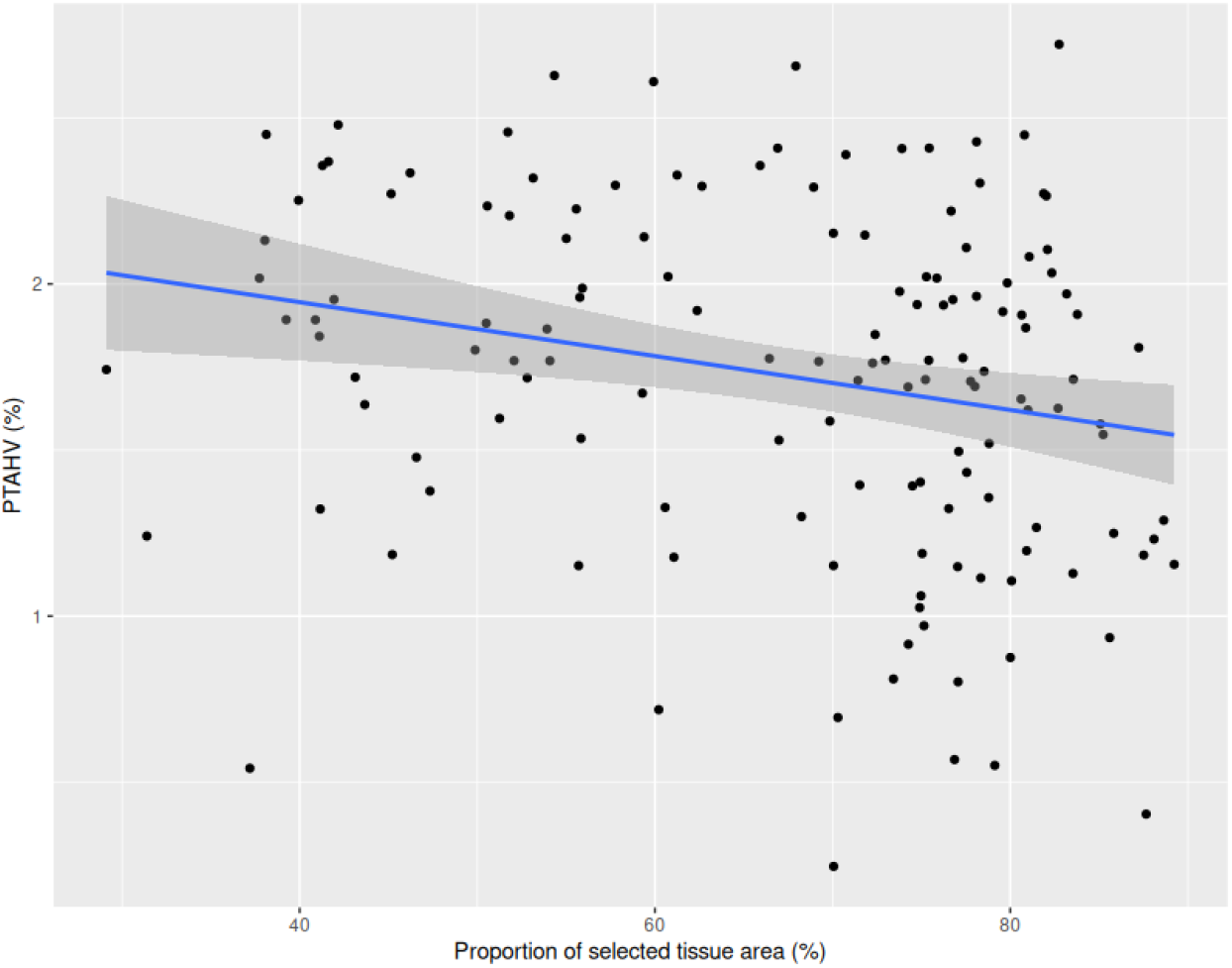
Plot showing the significant negative correlation (cor = -0.23, p < 0.05) between the proportion of selected tissue area (%) and PTAHV (%).

### Hepatocellular vacuolization

PTAHV ranges from 1.27% to 15.19% with an overall mean of 6.34% (± 3.02%). Significant differences in mean PTAHV (Anova type III, p < 0.05) were found between station and season. *P. flesus* from Seine has a higher mean PTAHV (6.88 ± 3.01%) compared to Canche (5.46 ± 2.86%). Fish sampled in Canche in winter display a mean PTAHV of 5.08% (± 3.48%) while fish sampled in Seine during summer rise up to 7.90% (± 3.03%). No significant difference (Anova type III, p > 0.05) was found in mean PTAHV between females and males (Figure 3).

**Figure 3.**
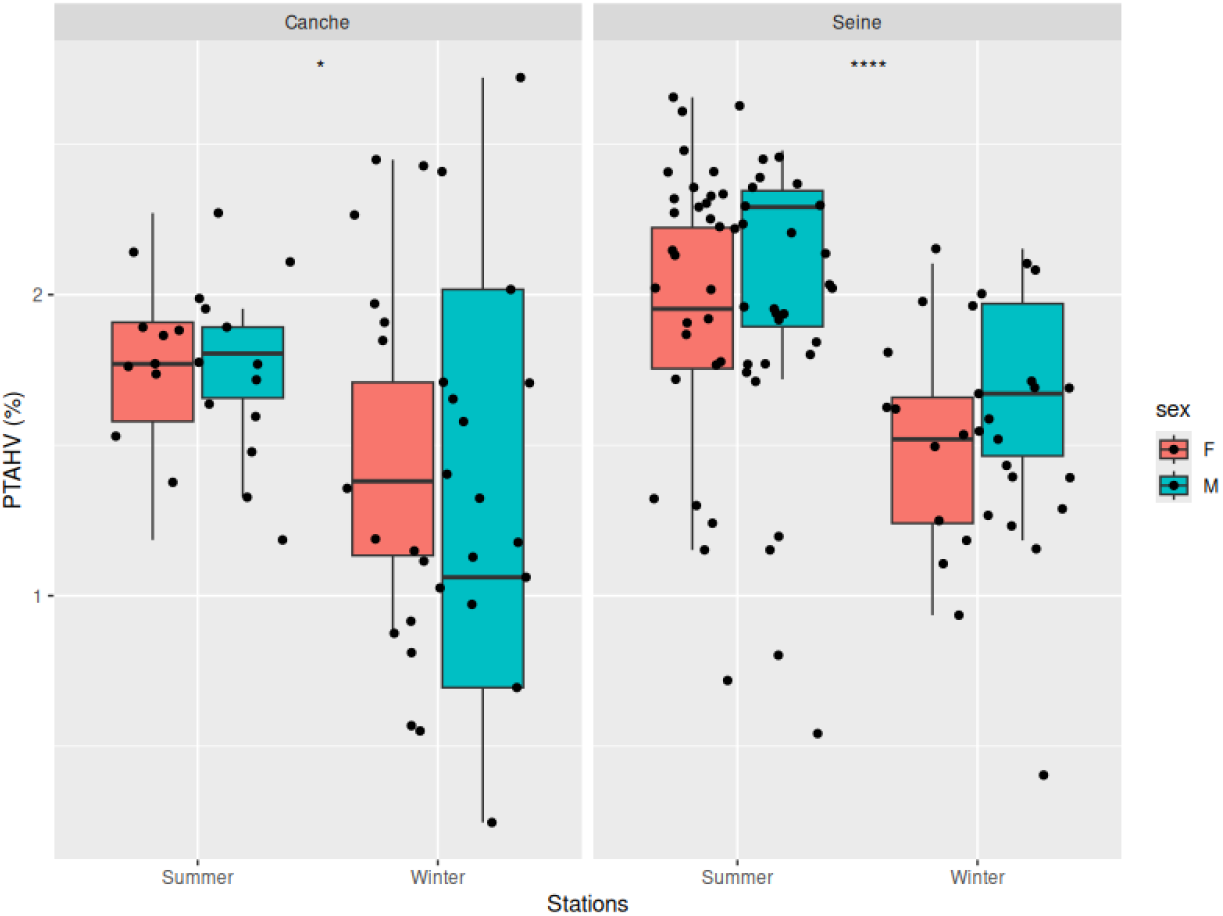
Box plot representing PTAHV in *P. flesus* according to station, season, and sex. Fish from Seine present a significantly higher proportion of hepatocellular vacuolization in summer (7.90 ± 3.03%) compared to winter (4.92 ± 1.76%), while there was no significant seasonal difference in fish from Canche. *P. flesus* from Seine having a higher mean PTAHV (6.88 ± 3.01%) compared to Canche (5.46 ± 2.86%).

### Fitness indexes

Fulton’s K index was negatively correlated with PTAHV (cor = -0.26, p < 0.05) (Figure 4.A). HSI was positively correlated with PTAHV (cor = 0.41, p < 0.05) (Figure 4.B). Age exhibits a significant cubic relationship with PTAHV (Figure 4. C).

**Figure 4.**
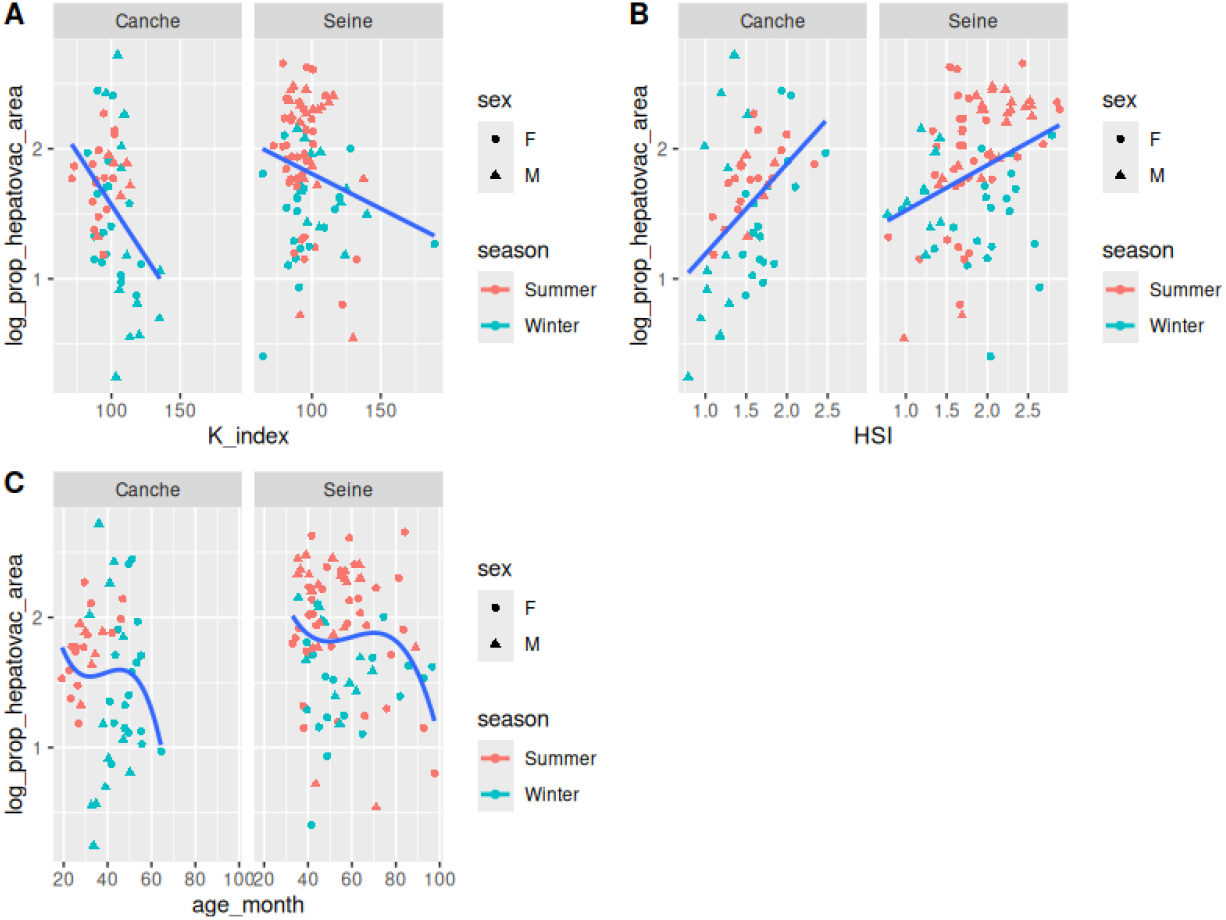
A) Plot representing the negative linear relationship (F = 10.54, p < 0.05, R2 = 0.06) and negative correlation (cor = -0.26, p < 0.05) between PTAHV and Fulton’s K index in *P. flesus* for the two stations. B) Plot representing the positive linear relationship (F = 28.89, p < 0.05, R2 = 0.16) and the positive correlation (cor = 0.41, p < 0.05) between PTAHV and HSI in *P. flesus* for the two stations. C) Plot representing the negative cubic polynomial relationship (F = 4.62, p < 0.05, R2 = 0.19) between PTAHV and age of *P. flesus* for the two stations.

### Reproductive markers

A significant negative correlation (cor = -0.3, p < 0.05) between PTAHV and E2 level measured in females was observed (Figure 5. A). A significant negative correlation (cor = -0.45, p < 0.05) was observed between the proportion of hepatocellular vacuolization and GSI for males *P. flesus* (Figure 5. B).

**Figure 5.**
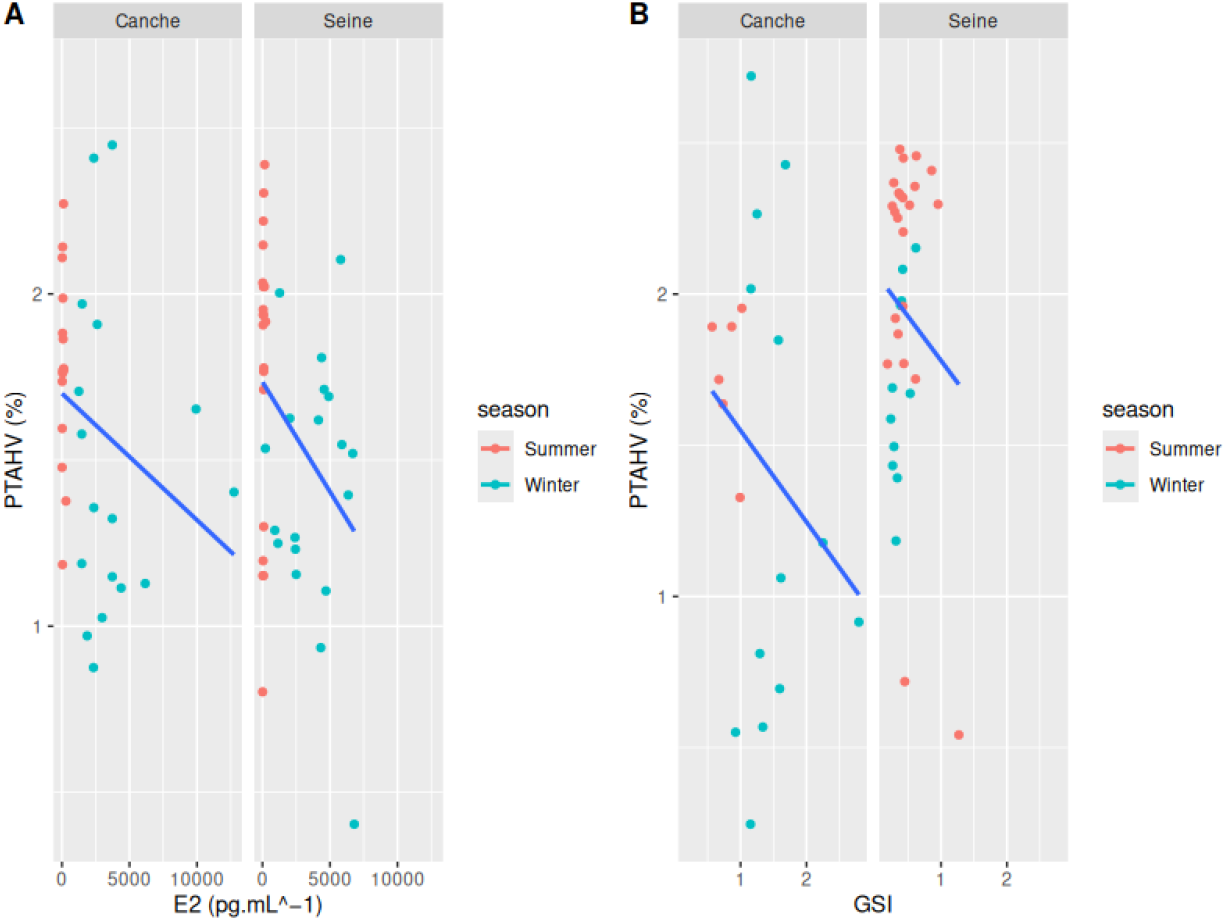
A) Plot representing the negative correlation between PTAHV and estradiol level (cor = -0.3, p < 0.05) measured on *P. flesus* females for the two stations. B) Plot representing the negative correlation between PTAHV and GSI (cor = -0.45, p < 0.05) measured on males *P. flesus* for the two stations.

### Liver histopathology

A significant difference (Anova type III, p < 0.05) was observed in the prevalence of liver histopathological categories between the different groups. *P. flesus* from Seine present a significantly higher prevalence of non-specific lesions and early toxicopathic non-neoplastic lesions compared to Canche. Other categories do not exhibit significant differences (Anova type III, p > 0.05) in their respective prevalence between station, season, and sex. The prevalence of nonspecific lesions shows a significant relationship with PTAHV (BRM estimates [95% CI] = 0.33 [0.18, 0.48]). Meaning that a 1-unit increase in non-specific lesion prevalence is associated with an increase of 0.33 units in the response variable. Other variables (early toxicopathic lesions, FCA) show no clear evidence of association with PTAHV.

A significant difference (Anova type III, p < 0.05) was observed in the prevalence of liver histopathological lesions between the different groups (Figure 6.A). *P. flesus* from Seine has a significantly higher prevalence of MMA, hydropic vacuolation and necrosis compared to individuals from Canche. A significant difference (Anova type III, p < 0.05) was observed in the prevalence of lipidosis between males and females, with males having a higher prevalence of lipidosis compared to females. Other lesions do not display a significant difference (Anova type III, p > 0.05) in their respective prevalence between station, season, and sex. The prevalence of lipidosis presents a significant positive relationship with PTAHV (BRM estimates [95% CI] = 0.48 [0.32, 0.64]). Meaning that a 1-unit increase in lipidosis prevalence is associated with an increase of 0.48 units in the response variable. Other variables (MMA, clear cell foci, phospholipidosis, granuloma abscess, necrosis) show no clear evidence of association with PTAHV (Figure 6.B).

**Figure 6.**
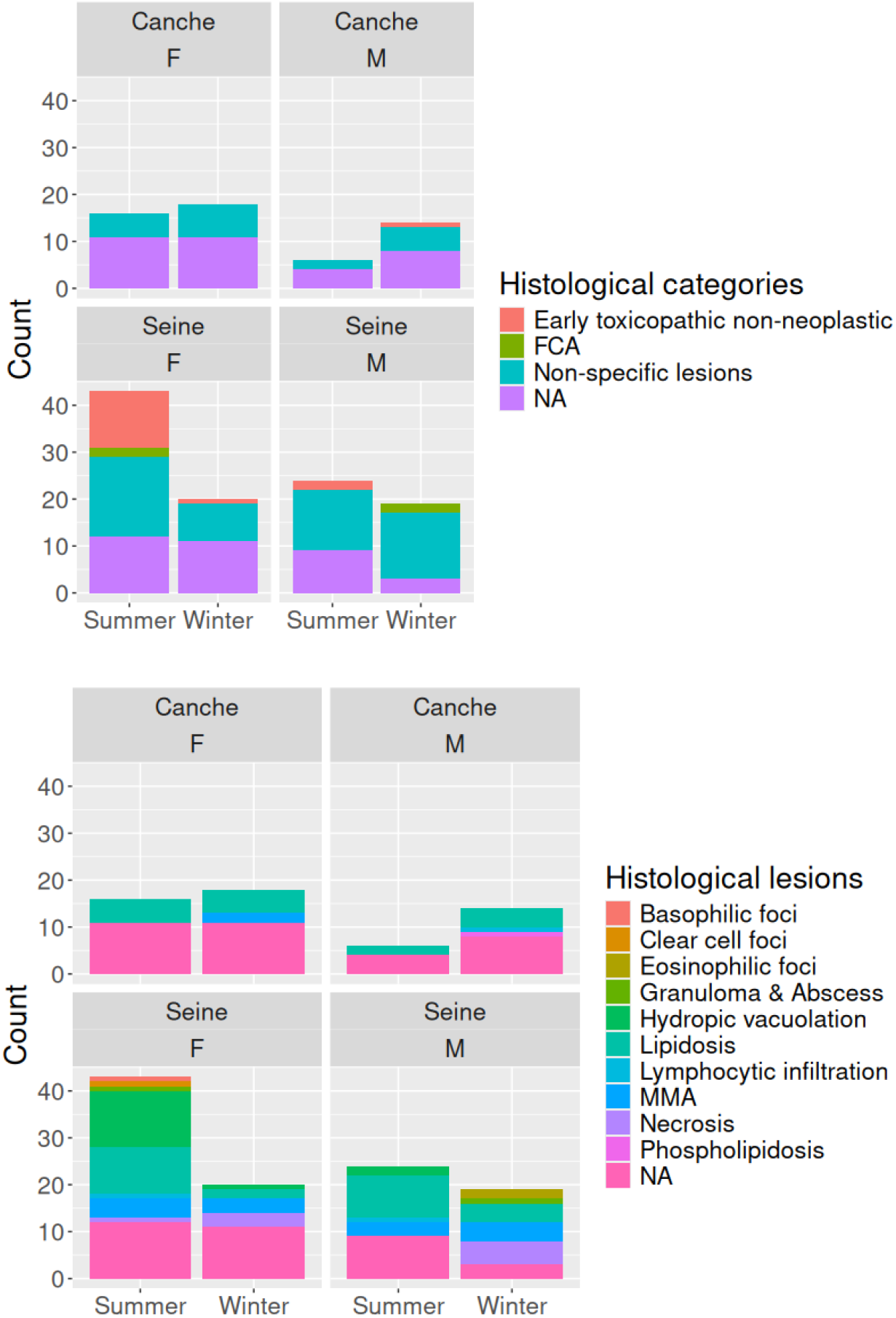
Liver histopathological categories’ count between station, season, and sex. Liver histopathological lesions’ count between station, season, and sex.

### Liver lipids content

A significant quadratic polynomial relationship between PTAHV and level of LPC was observed (Figure 7.A) and a cubic polynomial relationship between PTAHV and level of CL measured in the liver of *P. flesus* was observed (Figure 7.B). Significant negative correlation (cor = -0.33, p < 0.05) was observed between the level of total membrane lipids (*µ*g.mg^*−*1^) measured in *P. flesus* liver and PTAHV.

**Figure 7.**
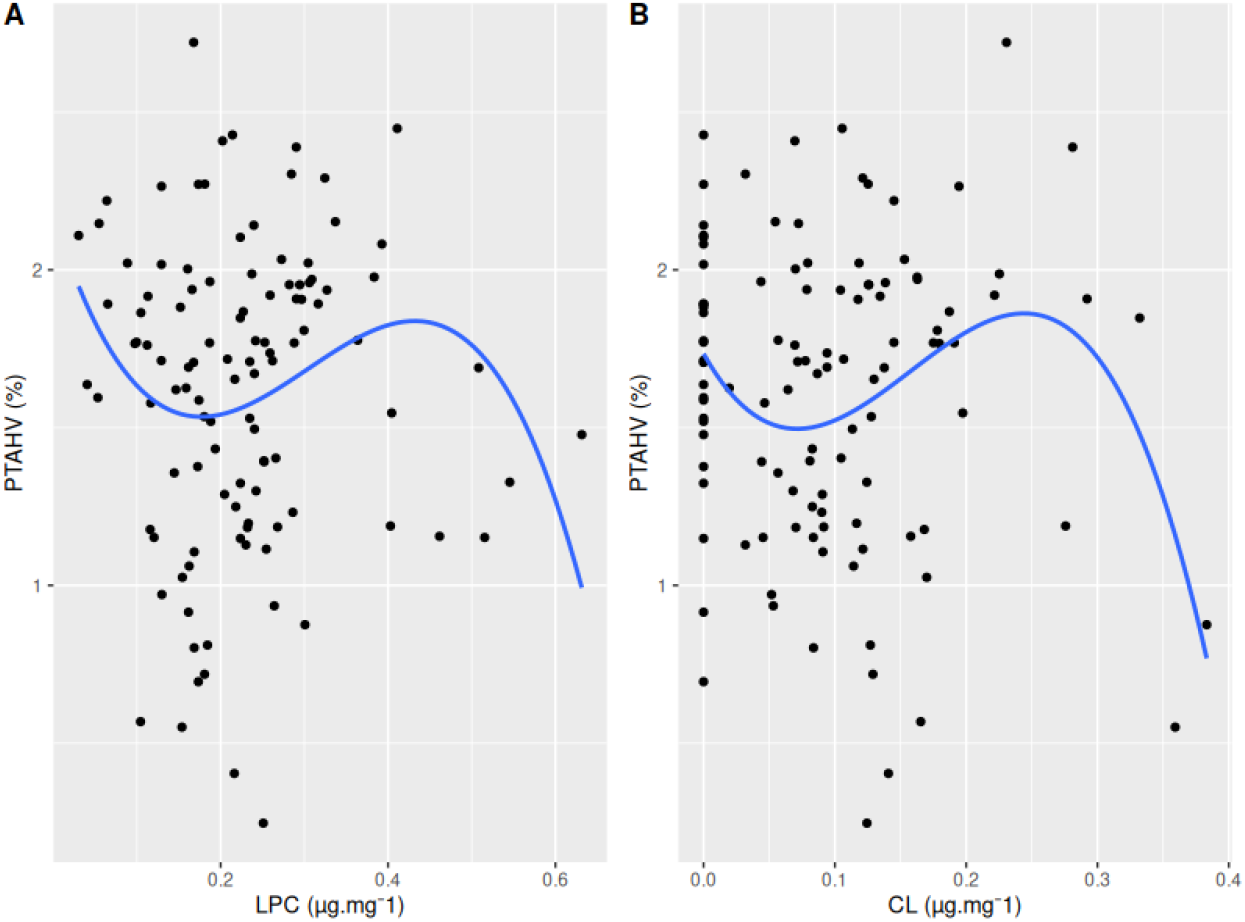
A) Plot representing the polynomial quadratic relationship between PTAHV and lipids LPC (*µ*g.mg^*−*1^) level measured in *P. flesus* liver. B) Plot representing the polynomial quadratic relationship between PTAHV and lipids CL (*µ*g.mg^*−*1^) level measured in *P. flesus* liver.

### Contaminants

#### Organic pollutants

- PCB A significant non-linear relationship between PTAHV and the levels of PCB 44, PCB 52, PCB 132 measured in the liver of *P. flesus* was observed (Figure 8).
- Pesticides A significant quadratic relationship was detected between PTAHV and the level of 4’4 DDE (*µ*g.kg^*−*1^) measured in *P. flesus* liver (Figure 9).
- Inorganic pollutants Significant positive quadratic relationship between PTAHV and level of Mn measured in the liver of *P. flesus*. Significant negative linear relationship between PTAHV and level of Cu, Hg, and Ni measured in the liver of *P. flesus* (Figure 10).

**Figure 8.**
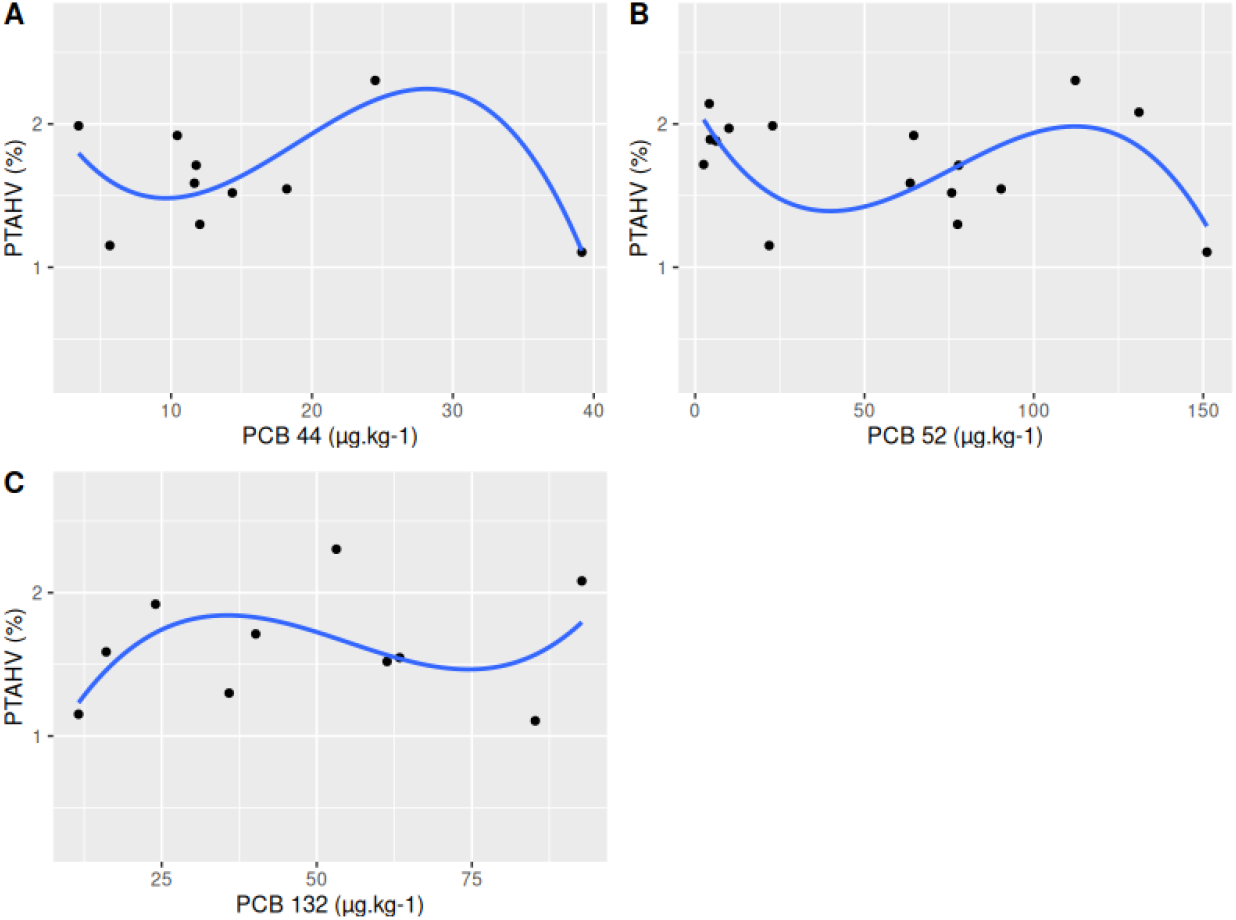
A) Plot representing the polynomial quadratic relationship between PTAHV and PCB 44 (*µ*g.kg^*−*1^) level measured in the liver of *P. flesus*. B) Plot representing the nonlinear relationship between PTAHV and PCB 52 (*µ*g.kg^*−*1^) level measured in the liver of *P. flesus*. C) Plot representing the polynomial cubic relationship between PTAHV and PCB 132 (*µ*g.kg^*−*1^) level measured in the liver of *P. flesus*.

**Figure 9.**
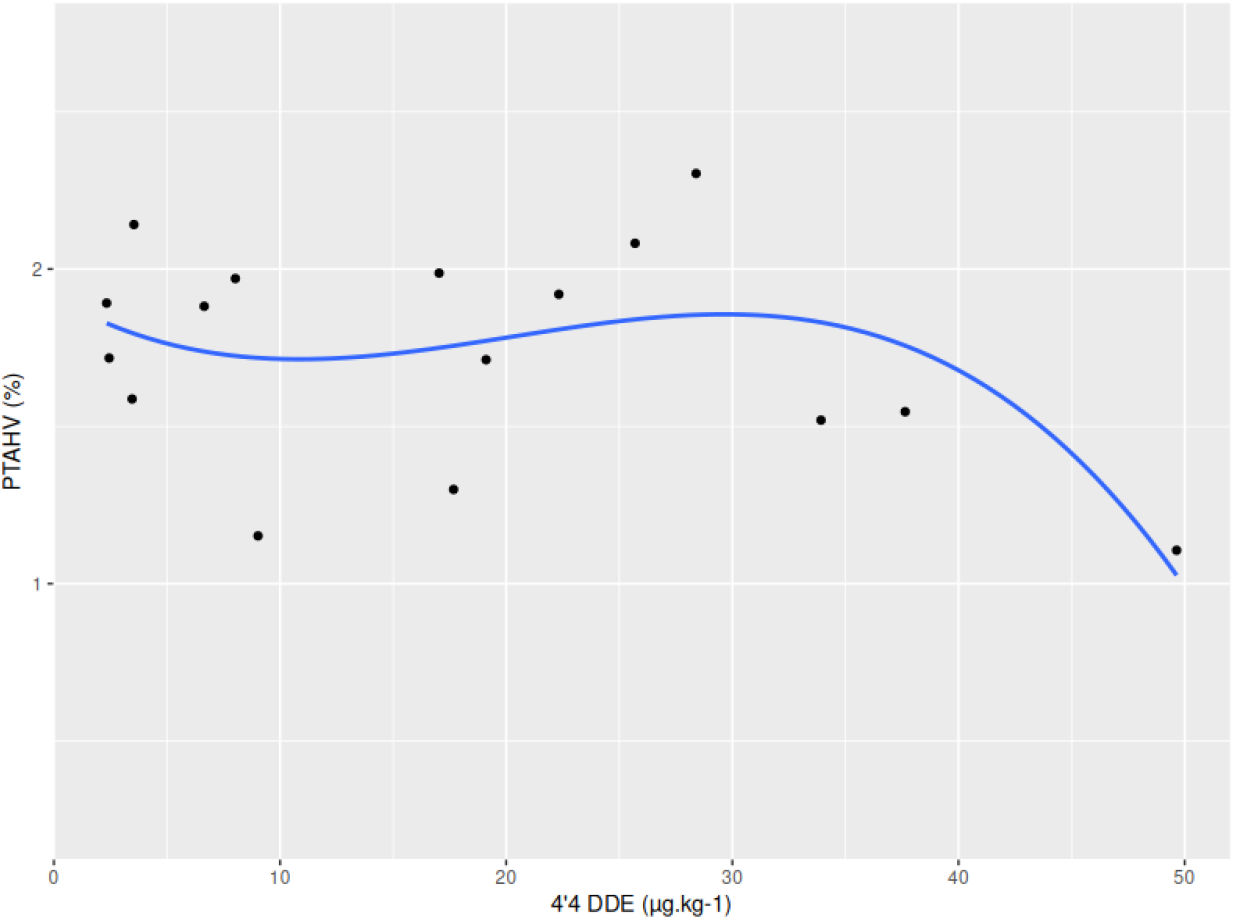
Plot representing the polynomial quadratic relationship between PTAHV and 4’4 DDE (*µ*g.kg^*−*1^) level measured in the liver of *P. flesus*.

**Figure 10.**
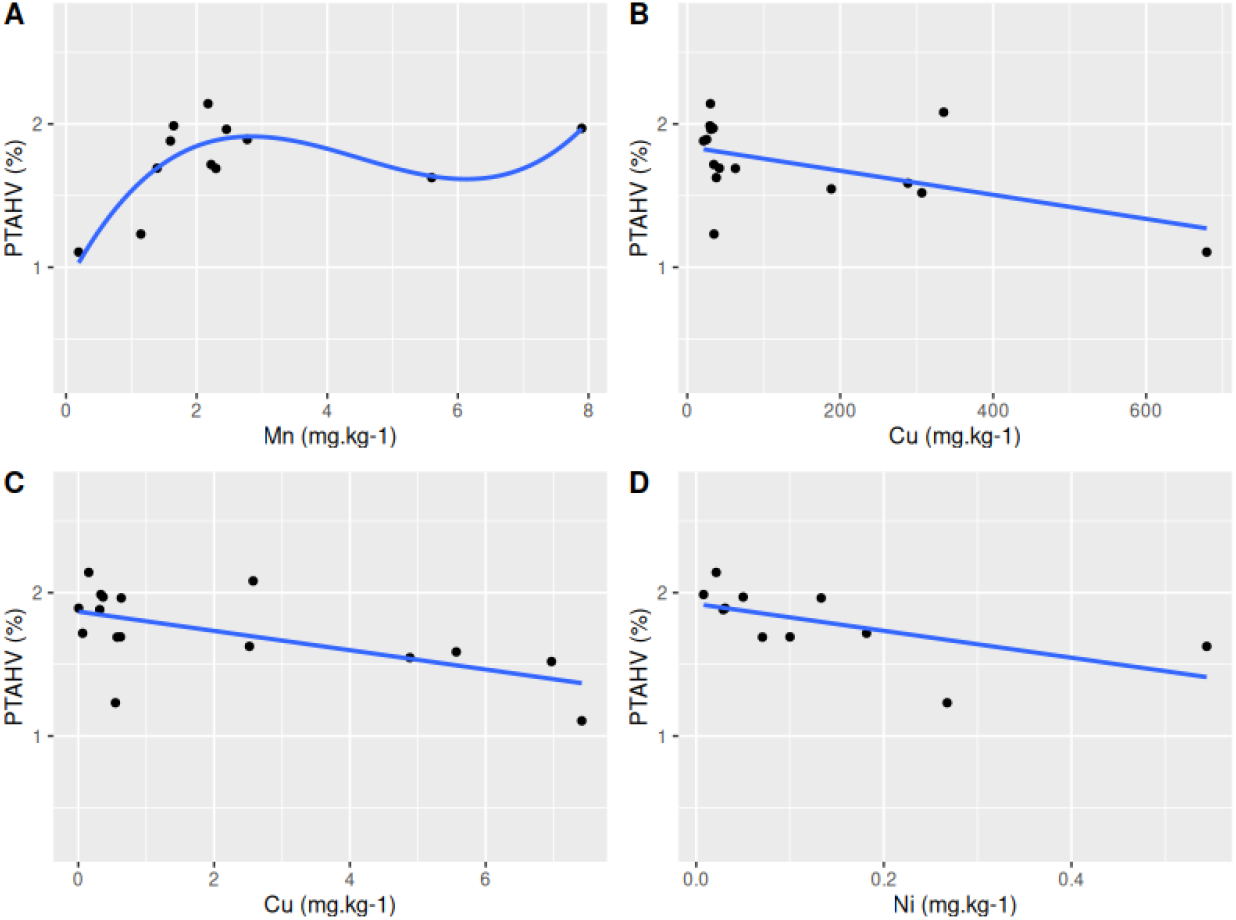
A) Plot representing the positive quadratic relationship between PTAHV and Mn (mg.kg^*−*1^) level measured in the liver of *P. flesus*. B) Plot representing the negative linear relationship between PTAHV and Cu (mg.kg^*−*1^) level measured in the liver of *P. flesus*. C) Plot representing the negative linear relationship between PTAHV and Hg (mg.kg^*−*1^) level measured in the liver of *P. flesus*. D) Plot representing the negative linear relationship between PTAHV and Ni (mg.kg^*−*1^) level measured in the liver of *P. flesus*.

### Mixture effect

A significant increase of PTAHV is associated with the overall increase of pollutant mixture in fish from the Seine estuary. After the 60th quantile, the confidence interval does not cross 0, meaning that the results were statistically significant. While PTAHV shows no significant variation with increased pollutant mixture for fish from Canche (Figure 11). Interaction plots show the really weak contribution of organic pollutants to PTAHV, while inorganic pollutant effects increase, and more specifically for Al, Cu, and Fe, with the increase of mixture concentration (Figure 12).

**Figure 11.**
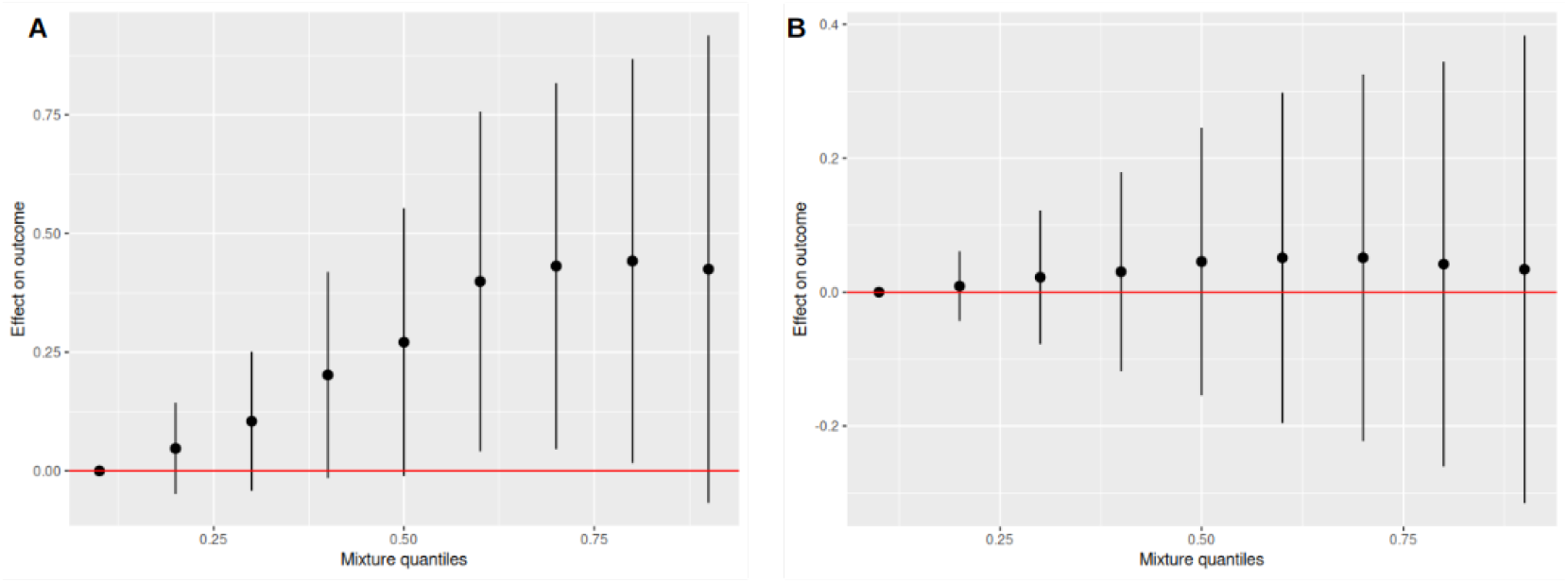
A) Plot presenting the overall effect of mixture quantiles on the response variable for fish from the Seine estuary. B) Plot presenting the overall effect of mixture quantiles on the response variable for fish from the Canche estuary.

**Figure 12.**
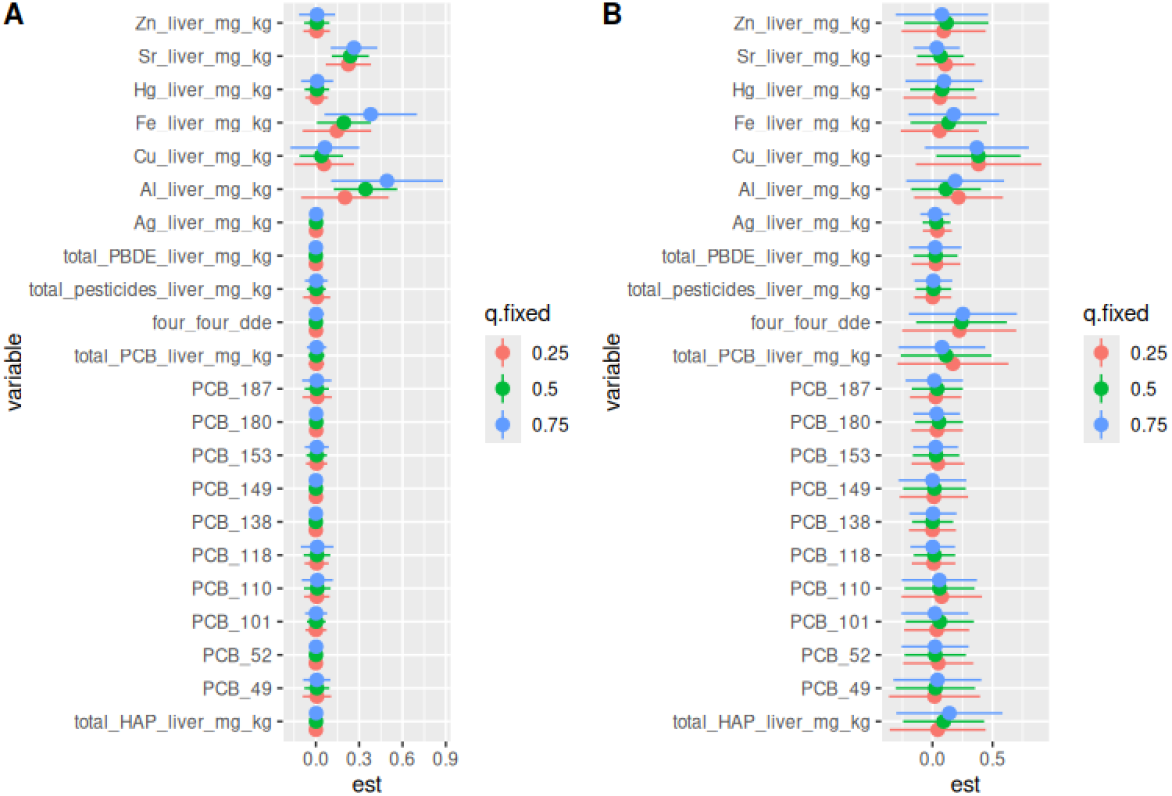
A) Interactions plot of mixture quantiles on the response variable for individuals from the Seine estuary. B)Interactions plot of mixture quantiles on the response variable for individuals from the Canche estuary.

## Discussion

In this article, a new WSI analysis method for the detection and quantification of hepatocellular vacuolization in flatfish liver is presented.

### Method validation

The difference in mean percentage of overlapping area was calculated by comparing manual annotation of area affected by hepatocellular vacuolization to the area detected by WSI analysis. The WSI analysis method matches at (72.5% ± 19.2%) the area defined by manual annotation. Method validation is crucial for the development and use of image analysis in order to measure its precision and reliability (Chen et al., 2021; Jain et al., 2024; Kong et al., 2011; Mulrane et al., 2008; Smith et al., 2021; Tizhoosh and Pantanowitz, 2018). A significant negative correlation was found between the proportion of selected tissue area and PTAHV. Various hypotheses are suggested to explain this negative relationship. The first hypothesis is that small liver samples tend to have a relatively smaller proportion of selected tissue area therefore, if the pathology is detected, the proportion of tissue impacted by hepatocellular vacuolization will be higher compared to bigger fish. Moreover, the tissue selection process is affected by the presence of artifacts in the tissue (tissue fracture, incision, puncture… ). These artifacts influence the proportion of selected tissue area as a similar artifact in terms of size, and will affect a relatively higher proportion of tissue in a small liver sample, as the entire tile will be removed from the analysis.

### Fitness indexes

Fish with a high condition index present a lower PTAHV, which is consistent with the fact that poor body condition is generally associated with diseases (Amara et al., 2007). The positive linear relationship observed between HSI and PTAHV could be explained by the fact that heavier livers were the ones accumulating more lipids and would exhibit more hepatocellular vacuolization. The negative relationship observed between age and PTAHV suggests that older fish would express this pathology less compared to younger fish (Stentiford et al., 2010). Fish populations from polluted areas might be more tolerant to xenobiotics compared to populations from less polluted areas. The tolerance can be expressed as physical adaptations, phenotypic plasticity, and selection of favorable genotypes across generations (Marchand et al., 2010, 2013).

### Reproduction marker

The negative linear relationship observed between E2 level and PTAHV, regardless of the station, could highlight the role of reserve lipids as a potential energy source in female’s reproductive cycle and more specifically for gonadal development (Dabrowska et al., 2017; Dando, 2011; Déniel, 1981; Gilliers et al., 2012; Kopko and Dabrowska, 2018). A negative linear relationship is observed between GSI and PTAHV on males, supporting the hypotheses of a potential depletion of hepatic reserve lipids during gonadal development (Kopko and Dabrowska, 2018).

### Liver histopathology

Histological analysis of liver damage has been extensively used as an integrated method for monitoring general biological effects and assessing the overall health status of fish, both at the individual and population levels. Inter-calibration and methodological standardization of histological analysis of liver pathologies in marine fish have enabled the application of this analysis in monitoring biological effects (Feist et al., 2004). Pathological signatures in flounder’s liver show a higher prevalence of histological symptoms in contaminated sites (Stentiford et al., 2010). Pathological signatures reflect the health status of individuals and are considered markers of major interest for better understanding the impact of chronic pollution on fish populations (Feist et al., 2004). The positive relationship between the prevalence of lipidosis and PTAHV was another evidence of the method’s accuracy, as results found by histopathological assessment were coherent with results found by WSI analysis.

### Liver lipids content

Neutral lipids are the primary form of lipid storage in fish. These reserves are mobilized during periods of high energy demand (growth, migration, fasting) as well as gonadal development during spawning seasons (Fonds et al., 1992). Exposure to pollutants could disrupt lipid metabolism, leading to lipid accumulation as a detoxification mechanism or due to impaired lipid catabolism (Barreto et al., 2018; Xie et al., 2026). However, no relationship was found between PTAHV and either neutral lipids or storage lipids levels. CL are phospholipids localized within the inner mitochondrial membrane, and are particularly rich in unsaturated fatty acids, which play a critical role for cellular energy production. CL are thus very susceptible to lipid peroxidation caused by oxygen radicals and other organic radicals (Hulbert et al., 2007). Its negative quadratic polynomial relationship with PTAHV could reflect mitochondrial dysfunction or adaptive responses to stress where energy production metabolism is reduced in case of contamination (Valencak and Azzu, 2014). LPC is a marker of membrane damage. Its quadratic polynomial relationship with the response variable may indicate that both low and high levels were associated with PTAHV, possibly due to membrane instability or oxidative stress (Borcier et al., 2020). The negative correlation between total membrane lipid level and PTAHV suggests that membrane lipid could be part of the lipid class accumulating in cytoplasmic vacuoles (Moore et al., 1997; Stehr et al., 1998). Membrane alteration by contaminant may trigger membrane lipid anabolism (Köhler et al., 2002).

### Contaminants and mixture effect

Fish contamination with persistent organic pollutants was significantly higher in the liver of

*P. flesus* from Seine vs. Canche. This could potentially be explained by the highly contaminated Seine sediments by POPs, relative to the sediments at the mouth of the Canche estuary (Laroche et al., 2013; Laurent et al., 2022). The negative quadratic relationship found between dioxin-like PCB congener (PCB 44, PCB 52) and PTAHV suggests that at high doses this type of PCB might interfere with lipid metabolism, leading to a decrease in hepatocellular vacuolization (Groh et al., 2022). On the other hand, the positive cubic relationship between chlorinated PCB congener (PCB 132) and PTAHV suggests that this contaminant, at high dose, would increase hepatocellular vacuolization in flounder liver (Ross, 2004). Associations between the incidence of some diseases in flatfish and the concentrations of PAHs in the sediment have been widely documented (Dupuy et al., 2014; Gilliers et al., 2012; Lyons et al., 2004). Generally, most PAHs are efficiently metabolized in fish tissues, and measuring metabolites in the bile is usually preferred (Thain et al., 2008). The effect of pesticides on *P. flesus* has been documented to alter metabolic functions, including energy production, general metabolism, signaling, transport, immune system, and structure (Evrard et al., 2010; Marchand et al., 2006). The quadratic polynomial relationship between 4’4 DDE level and PTAHV suggests that at high concentration, this pesticide would decrease hepatocellular vacuolization. Some pollutants are metabolized because of their shorter half-lives, others, such as heavy metals, pesticides, persistent organic pollutants, and other endocrine-disrupting chemicals, continue to accumulate in tissues and have significant and long-term effects on organisms’ fitness (Liu and Sayes, 2024; Zhu et al., 2024). The negative correlation observed between PTAHV and the level of Cu, Hg and Ni measured in the liver of Seine *P. flesus* highlight potential inhibitory effect of these metals on lipid metabolism. However, most studies focus on single pollutant’s effect and do not accurately reflect realistic conditions since organisms are exposed to a combination of several xenobiotic substances at any given time, which might have antagonistic or synergistic effects. The additive effect in a multipollutant mixture refers to the assumption that the combined health impact of multiple pollutants is equal to the sum of their individual effects. This approach is commonly used in environmental studies, although it may not always accurately reflect the complex interactions between different pollutants (Rodea-Palomares et al., 2015; Zhu et al., 2024). As the results of the mixture effect analysis show, the more the whole mixture increases in concentration, the greater the effect seen on the response variable for fish from the Seine estuary. Metal effects seem to increase with mixture concentration increase. Therefore, we can speculate that inorganic pollutants may have some sort of interaction with other compounds in the mixture. On the opposite, the response variable does not significantly vary when the whole mixture increases in fish from the Canche estuary. These results suggest that the PTAHV rate may be majorized in a multipollutant context, which is another layer of complexity to account for when using this pathology as a histological biomarker.

## Conclusion

In the general context of global change, marine organisms are increasingly subjected to multistress which in turn increases their susceptibility to pollution (Lapointe et al., 2011). Toxicopathic liver lesions in fish are playing an important role as biomarkers of contaminant effects in marine biological monitoring programs, and therefore the use of histological assessment is essential in order to distinguish non-specific lesions to pathological changes due to xenobiotic contaminants (Wolf and Wheeler, 2018). In addition, because there is evidence of a general decline in the overall prevalence of macroscopically visible hepatic neoplasms in some flatfish species from Europe caused by a general diminution of pollutants in European rivers, it is likely that greater attention will be given to the prevalence and extent of microscopic lesions in livers exhibiting no evidence of gross lesions (Lang et al., 2017). Moreover, the fish’s ability to survive extensive liver necrosis suggests they may be valuable animal models for studying sublethal hepatic toxicity and liver regeneration (Wolf and Wolfe, 2005). For this, there is a pressing need for the standardization and development of more precise methods of histopathological diagnosis (Feist et al., 2004). Exploring the effects of polluting cocktails in natural conditions, in particular by developing a collaborative approach between work in human health and animal models in ecosystems, is also needed (Barouki et al., 2023). In the future, a comparative fish-human pathology approach could be adopted, knowing that fish models are an appropriate alternative to mammalian models in toxicity and pathology studies (Gonçalves et al., 2020; Sepp and Giraudeau, 2023).

## Acknowledgements

Thanks to Silja Kavlie Fykse from the Department of Pathology at Stavanger University Hospital for scanning the slides. Thanks to Nicolas Jovanovic from Grenoble Alpes University for his help with the bkmr analysis.

## Data, script, code, and supplementary information availability

Original data came from Laurent et al., 2022 (https://doi.org/10.1007/s11356-022-20000-y). Dataset available on the following repository: https://doi.org/10.5281/zenodo.23457131. Script and codes are available online (https://github.com/Valentin-Geslin/hepatovac_fish_liver).

